# Specific enrichment of hyperthermophilic electroactive *Archaea* from deep-sea hydrothermal vent on electrically conductive support

**DOI:** 10.1101/272039

**Authors:** Guillaume Pillot, Eléonore Frouin, Emilie Pasero, Anne Godfroy, Yannick Combet-Blanc, Sylvain Davidson, Pierre-Pol Liebgott

## Abstract

While more and more investigations are done to isolate hyperthermophilic exoelectrogenic communities from environments, none have been performed yet on deep-sea hydrothermal vent. Samples of black smoker chimney from Rainbow site on the Atlantic mid-oceanic ridge have been harvested for enriching exoelectrogens in microbial electrolysis cells under hyperthermophilic (80°C) condition. Two enrichments have been performed: one from direct inoculation of crushed chimney and the other one from inoculation of a pre-cultivation on iron (III) oxide. In both experiments, a current production was observed from 2.4 A/m^2^ to 5.8 A/m^2^ with a set anode potential of +0.05 vs SHE. Taxonomic affiliation of the exoelectrogen communities obtained exhibited a specific enrichment of *Archaea* from *Thermococcales* and *Archeoglobales* orders on the electrode, even when both inocula were dominated by *Bacteria*.

## 1. Introduction

Since the discovery of the first deep-sea hydrothermal vent in 1977, many studies have expanded our understanding of extremophilic life forms in those environments. The “black smoker” deep-sea hydrothermal vents, located along the ridges of the Atlantic, Pacific and Indian oceans, are the result of volcanic activities that generate hydrothermal chimneys composed of polymetallic sulfide minerals (heterogeneous pyrite) (Dick et al. 2013). Those environments exhibit dynamic habitats that are characterized by large steep thermal and chemical gradients. These gradients provide a wide range of growth conditions for many extremophilic microorganisms growing as biofilms and being the base of these specific deep-sea ecosystems (Flores et al. 2011; Kristall et al. 2006). In these environments has been shown the presence of thermophilic microorganisms from the *Archaea* domain, mainly belonging to the orders *Thermococcales*, *Methanococcales*, and *Archaeoglobales*, whereas mesophilic and thermophilic microorganisms from the *Bacteria* domain belong to *Epsilon-proteobacteria* (Huber et al. 2010; Vetriani et al. 2014) and to the orders *Aquificales* and *Thermotogales* (Miroshnichenko and Bonch-Osmolovskaya, 2006). Considering the mineralogical composition of these hydrothermal chimneys (polymetallic massive sulfide), it appears likely that a significant proportion of its microbial populations is dependent on energetic metabolisms based on the dissimilatory reduction of insoluble “metals” or sulfurs compounds (Cao et al. 2014; Konn et al. 2015).

As the microbial cell envelope is neither physically permeable to insoluble minerals nor electrically conductive, microorganisms have evolved strategically to exchange electrons with insoluble extracellular minerals, a mechanism known as the extracellular electrons transfer (EET) (Hinks et al. 2017; Shi et al. 2016). The first described EET capable bacteria were *Shewanella* and *Geobacter* (Shi et al. 2007). The role of membrane bound electron transport chains in carrying out EET was well acknowledged, but the exact mechanisms are still not completely understood (Kumar et al. 2017). Most of the knowledge about EET in microbes is derived from studies in bioelectrochemical systems (Allen and Bennetto, 1993; Logan et al. 2006; Schröder et al. 2015) which helped to define a novel group of microorganisms called exoelectrogens. These exoelectrogenic microorganisms are capable of extracellular electron transfer to a solid electrode, which are used in Microbial Electrochemical Technologies (METs) such as Microbial Fuel Cells (MFC) and microbial electrolysis cells (MEC) (Doyle and Marsili, 2015). In the past years, exoelectrogenic activity has been reported in almost 100 microbial species which are mostly affiliated with the bacterial phylum *Proteobacteria* (Koch and Harnisch, 2016). It includes the extensively studied *Geobacter sp*. (Busalmen et al. 2008; Reguera et al. 2006) and *Shewanella sp*. (Gorby et al. 2006; Marsili et al. 2008), as well as *Cyanobacteria* (J. McCormick et al. 2011; Sekar et al. 2014). All these microorganisms are mesophilic and grow optimally at moderate temperatures, ranging from 20°C to 45°C. To date, only four thermophiles, *Thermincola ferriacetica* (Marshall and May, 2009), *Thermincola potens* strain JR (Wrighton et al. 2011), *Calditerrivibrio nitroreducens* (Fu et al. 2013) and *Thermoanaerobacter pseudethanolicus* (Lusk et al. 2015), all isolated from extreme natural environments, were shown to generate electricity in MFCs operating at a temperature higher than 50°C. Moreover, it is only very recently that two hyperthermophilic strains, *Pyrococcus furiosus* and *Geoglobus ahangari,* have been shown to have the capacity to produce electricity in MFC or MEC (Sekar et al. 2017; Yilmazel et al. 2018). This would be the two first hyperthermophilic archaeon strains described having the capacity of EET. Several studies reported the enrichment of mixed cultures of efficient thermophilic exoelectrogens on the anode of METs (Ha et al. 2012; Jong et al. 2006; Mathis et al. 2007; Wrighton et al. 2008). More recently, two exoelectrogenic biofilms have been enriched at 70°C from a high-temperature petroleum reservoir (Fu et al. 2015) and at 80°C from Red Sea brine pools (Shehab et al. 2017) on the anode of MFC systems. To the best of our knowledge, no exoelectrogenic biodiversity enrichment from deep-sea hydrothermal vents has been achieved yet, while it seems likely that some microbial populations require this type of energy metabolism to expand in these particular environments. Interestingly, a MFC was installed at a hydrothermal vent field (Girguis and Holden, 2012) showing for the first time, the *in situ* electricity generation in those extreme ecosystems. More recently it has been demonstrated that there is a widespread and distant electron transfer through the electrically conductive hydrothermal chimney by the internal oxidation of the hydrothermal fluid coupled to the reduction of oxygenated seawater at the external side of the chimney (Yamamoto et al. 2017). This electricity generation in deep-sea hydrothermal systems must affect the surrounding biogeochemical processes and the development of microbial communities through potential EET-capable microorganisms.

The aim of this study was to promote and identify a part of the exoelectrogenic microbial community from a hydrothermal chimney of the Rainbow site on the Atlantic mid-oceanic ridge. For this investigation, two experiments were carried out using a semiautomatic two-chamber BioElectrochemical System Stirred-Reactor (BES-SR) prototype (graphical abstract), operating at high temperatures (80°C). The first one was performed to enrich microbes in our BES-SR system, directly from a chimney fragment. The second enrichment in BES was carried out from an inoculum obtained by a microbial enrichment culture in flask on Fe(III) oxide particles as electron acceptor using the same hydrothermal sample. This was done in order to pre-cultivate potential exoelectrogenic microbes on iron oxide and observing the impact on biodiversity obtained subsequently in BES. The evolution of current production, of microbial diversity and biomass during the different enrichments have been studied through electrochemical method and molecular biology survey.

## 2. Material and methods

### 2.1. Sample collection and preparation

The inoculum used for all the experiments was collected on the Rainbow hydrothermal vent field (36°14’N, MAR) by the Remote Operated Vehicle VICTOR 6000 during EXOMAR cruise in 2005 led by IFREMER (France) on board RV L’Atalante (Godfroy, 2005). Sample (EXO8E1) was collected by breaking off a piece of a high temperature active black smoker using the arm of the submersible and bring back to the surface into a decontaminated insulated box. On board, chimney fragments were anaerobically crushed into an anaerobic chamber (La Calhene, France) and stored into flasks under anaerobic conditions (anoxic seawater at pH 7 with hydrogen sulfide and N2:H2:CO2 (90:5:5) gas atmosphere) and kept at 4°C. Prior to the experiment, pieces of hydrothermal chimney were removed from the sulfidic seawater flask, crushed in a sterile mortar and pestle in an anaerobic chamber (Coy Laboratories Inc.) and distributed in anaerobic tubes for further different experiments.

### 2.2. Enrichment on iron oxide in flask

To obtain enrichment of electro-active microbes on insoluble iron (III) oxide as electron acceptor, 2.5g of crushed hydrothermal chimney was inserted in 500 ml flasks under nitrogen gas atmosphere filled with 250ml of mineral medium at pH 7 containing 30 g/l NaCl, 0.65 g/l KCl, 0.5 g/l NH_4_Cl, 0.3 g/l KH_2_PO_4_, 0.3 g/l K_2_HPO_4_, 0.1 g/l MgCl_2_, 0.1 g/l CaCl_2_, 0.35 g/l Cysteine HCl,
0.2 g/l of yeast extract and 10ml/l of Balch trace mineral solution (Uzarraga et al. 2011). The media was supplemented with 1g/l of iron (III) oxide, and 10mM of acetate or 4g/l of yeast extract as electron donor and carbon source. The flasks were incubated at 80°C in static condition until a dark coloration of iron oxide was observed. Two subcultures were performed in the same condition with 1% of inoculum from the previous culture.

### 2.3. Semi-Automated Bioelectrochemical Systems

A prototype of semi-automated BioElectrochemical System in Stirred-Reactor (BES-SR) has been developed to assess the enrichment of hyperthermophilic electroactive microorganisms. The system was composed of a two-chamber jacketed glass reactor (Verre Labo Mula, France) with a 1.5L working volume, thermostated with a Heating Circulator (Julabo SE 6, France) at 80°C ± 1 °C, and separated by an Anion Exchange membrane (Membrane International Inc.). The working electrode was a 20 cm^2^ carbon cloth (PaxiTech SAS, France) with a 3M Ag/AgCl reference electrode and the counter electrode a 20cm^2^ carbon cloth coated with platinum (Hogarth, 1995).

The bioelectrochemical system was connected to a semi-automated platform previously described (Boileau et al. 2016) to control the composition and rate of gas input (H_2_, CO_2_, O_2_, N_2_) with mass flowmeters (Bronkhorst, Netherlands) and a continuous monitoring of output gas composition (H_2_, N_2_, CH_4_, O_2_) using a micro-GC equipped with a catharometric detector (MS5A, SRA Instrument, France). To ensure anaerobic condition, the culture medium was continually sparged with a 50 ml/min flow of N_2_. The pH was maintained at 7 ± 0.1 by the addition of sodium hydroxide (NaOH 0.5 mmol/L) or hydrochloric acid (HCl 0.5 mmol/L). The stirring, driven by two axial impellers, was set to 150 rpm. The measurements of pH and ORP (Mettler Toledo InPro 3253, Switzerland), temperature (Prosensor pt 100, France), stirring, CO_2_ with a CARBOCAP CO2 probe (Vaisala GMT 221, Finland), H_2_, N_2_, CH_4_, O_2_, bioreactor liquid volume, NaOH and HCl consumption were measured and managed by the BatchPro software (Decobecq Automatismes, France). SP-240 potentiostats and EC-Lab software (BioLogic, France) were used to poise the electrodes at a fixed potential and measure current. All the measured currents are expressed in ampere per square meter of electrode area.

### 2.4. Operating conditions in BES-SR

Before each experiment, the bioelectrochemical system was dismantled, washed, and sterilized by autoclaving at 120 °C for 20 min. Then, the system was connected to the platform and 1,5L of mineral medium was injected and supplemented with 10 mM acetate and 0.15 g/l of Yeast Extract (YE). The liquid media was set in the operational condition for a few hours prior to perform a cyclic voltammetry (CV, 20 mV/s). First, the system (BES1) was inoculated with 15 g of crushed hydrothermal chimney in anaerobic condition. A chronoamperometry was carried out by the potentiostat to poise the electrode at +0.05V vs Standard Hydrogen Electrode (SHE), and a measurement of the current was taken every 10s. A cyclic voltammetry was also performed at the end of the experiment. Secondly, a new system (BES2) was inoculated in the same condition with 1% of the last enrichments in flask on iron (III) oxide on yeast extract. An abiotic control and an inoculated but non-polarized control have been performed in the same conditions to exhibit the exoelectrogenic specificity of the biofilm on the polarized electrode.

### 2.5. Taxonomic and phylogenetic classification

Taxonomic affiliation was performed according to (Zhang et al. 2016). DNA were extracted from 1g of the crushed chimney and at the end of each experiment: from 1g of liquid from flask enrichment, on the scrapings from half of the 20cm^2^ of working electrode and from 50ml of the liquid media of the BES and flask enrichments, which were centrifuged and suspended in 1ml of sterile water. The DNA extraction was carried out using the MoBio PowerSoil DNA isolation kit (Carlsbad, CA USA). The V4 region of the 16S rRNA gene was amplified using the universal primers 515F (5′-GTG CCA GCM GCC GCG GTA A-3′) and 806R (5′-GGA CTA CNN GGG TAT CTA AT-3′) with Taq&Load MasterMix (Promega) and PCR reactions were carried out using C1000 Thermal Cycler (BioRad) with the following conditions: initial denaturation at 94°C for 3 min followed by 30 cycles of denaturation at 94°C for 30 s, primers annealing at 50°C for 30 s and extension at 72°C for 90 s, followed by a final extension step of 5min at 72°C. The amplified gene regions were sequenced on Illumina MiSeq 2500 platform (GeT-PlaGe, France) to generate paired-end 150bp reads. The reads were merged using the FLASH software. The taxonomic affiliation was performed with the QIIME software package v 1.9.1. Chimera were removed from the merged sequenced using UCHIME Algorithm. Then, the filtered sequences were clustered into OTUs using the RDP method with a minimum bootstrap confidence of 0.8 and OTUs were affiliated using the Silva database as reference. To analyze the alpha diversity, the OTU tables were rarified to a sampling depth of 10770 sequences per library and two metrics were calculated: the richness component represented by the number of OTUs observed, and the Pielou’s index representing the evenness component of our biodiversity. The construction of clone library was performed by amplifying the 16S rRNA genes by previously described PCR method with a FD1 (5’-AGAGTTTGATCCTGGCTCAG-3’) and R6 (5’-TACGGCTACCTTGTTACG-3’) primer set using Taq&Load MasterMix (Promega). The amplicons were cloned into the pGEM-T easy vector (Promega) and transformed into *E. coli* JM109 competent cells that were grown overnight in LB agar at 37°C. Fragment of 16S rRNA gene (~1300 bp) of the clones were sequenced by ABI3730xl (GATC Biotech) sequencers.

### 2.6. Quantitative PCR of archaeal and bacterial 16S rRNA gene copies

Bacterial and archaeal quantification was carried out by qPCR with SsoAdvanced™ Sybr Green Supermix on a CFX96 Real-Time System (C1000 Thermal Cycler, Bio-Rad Laboratories, CA, USA) with the primers DGGE300F (5’-GCC TAC GGG AGG CAG CAG-3’) and Univ516 (5’- GTD TTA CCG CGG CKG CTG RCA-3’) specific to Bacteria and Arc931F (5’-AGG AAT TGG CGG GGG AGC A-3’) and m1100R (5’-BTG GGT CTC GCT CGT TRC C-3’) for Archaea. The PCR program was composed of a 10s denaturation step at 94°C, a hybridization step of 10s at 55°C (Bacteria) or 62°C (Archaea) and a 10s elongation step at 72°C, with melting curves performed at the end of each reaction to ensure product specificity. A standard curve from 10^2^ to 10^10^ 16S rRNA gene copies was obtained by diluting pGEM-T plasmids harboring hyperthermophilic bacterial or archaeal rRNA gene fragment obtained from microbial community of interest. The results were expressed in copies number of 16s rRNA gene per gram of crushed chimney, per milliliter of liquid media or per cm^2^ of surface of electrode.

## 3. Results and discussion

### 3.1. Microbial diversity of hydrothermal chimney from Rainbow site of Atlantic Ocean

Prior to study specifically the electroactive community putatively present on a chimney of the hydrothermal Rainbow site, an analysis of the total microbial diversity present in our crushed chimney inoculum was done by using the Illumina method. The chimney biodiversity (figure 1) was composed of 160 OTUs representing a high richness of species. As indicated by the Equitability index (0.677), the sequences were distributed relatively equally in the different OTUs. Furthermore, the taxonomic affiliation of the 160 OTUs showed that 66% were assigned to *Bacteria* and 33% to *Archaea* domains. This was consistent with the quantitative Polymerase Chain Reaction (qPCR) assays which showed a dominance of *Bacteria* (figure 2) compared to *Archaea* (6.38 ± 0.05 vs 4.77 ± 0.19 log of 16S rRNA gene copies per gram of chimney, respectively). The difference of ratio obtained can be explained by the bacterial quantification by 16S rRNA gene qPCR, always overestimated because of the high number of gene copies per cells compared to *Archaea* where the number of 16S rRNA gene copies is generally equal to 1 per cell (cf. rrnDB, (Stoddard et al. 2015)). Within the bacteria domain, the OTUs were mainly assigned to 5 phyla: *Proteobacteria* (58%), *Firmicutes* (3%), *Aquificae* (1%), *Bacteroidetes* (1%) and *Thermodesulfobacteria* (1%) whereas the archaeal OTUs were only identified as *Euryarchaeota* represented by *Thermococcales* (23%) and *Archaeoglobales* (11%). This taxonomic profile was substantially similar to those previously reported on Rainbow chimneys (Cerqueira et al. 2017; Flores et al. 2011), except for the proportion of methanogens and *Desulfurococcales* found significantly lower in our samples. Despite a recognized stability in the conservation of hyperthermophilic microorganisms at low temperature (Wirth, 2017), our samples have been conserved more than 12 years at 4°C, this could mostly explain the decrease in Archaea diversity.

**Figure 1:**
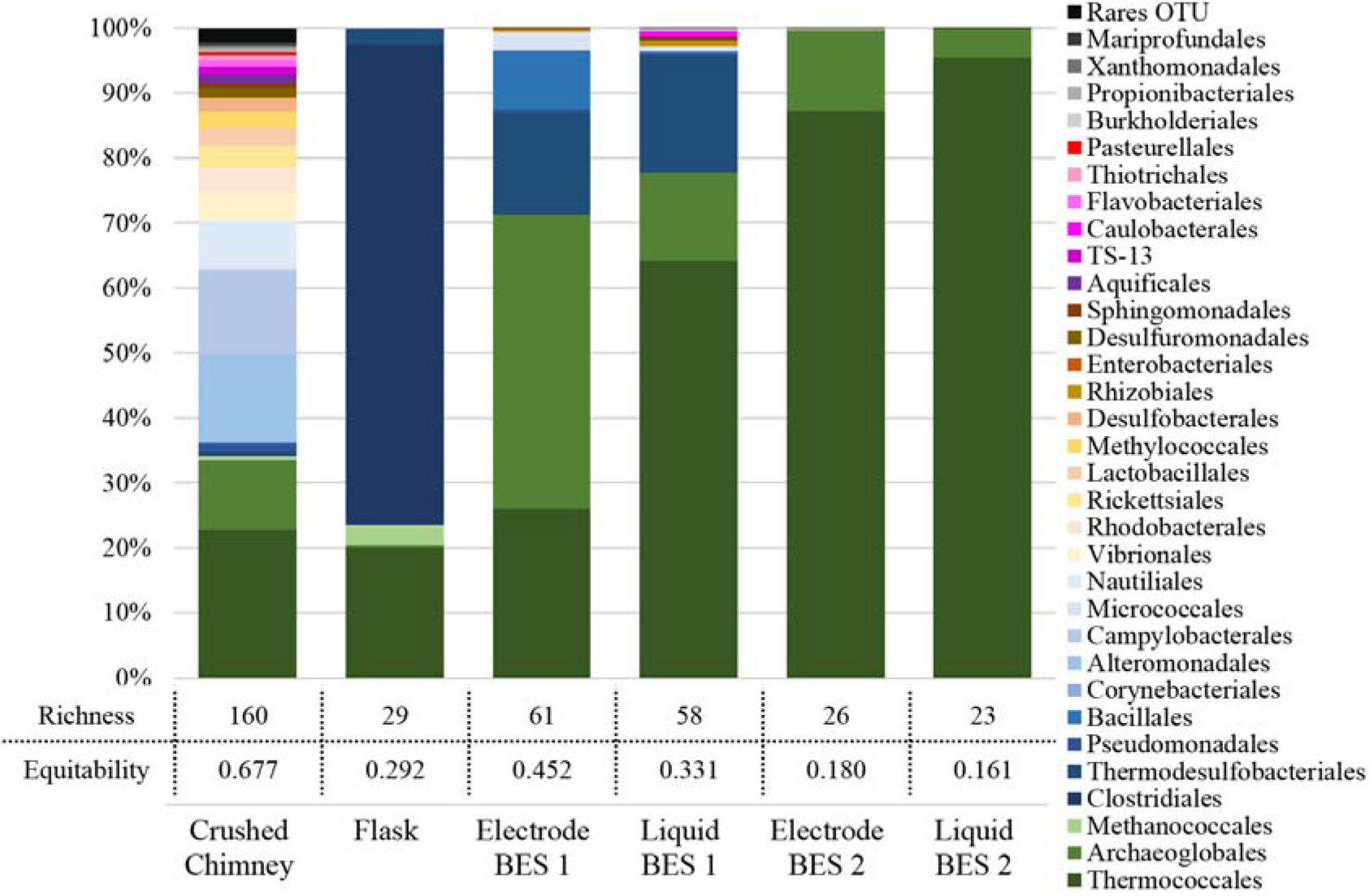
Dominant taxonomic affiliation and biodiversity indices of microbial communities from Crushed Chimney, Flask enrichment on iron (III) oxide, Electrode and Liquid media from BES1 and BES2. OTUs representing lower than 0.5% of total sequences of the sample was grouped as Rares OTU. Biodiversity indexes of Richness and Equitability represent the number of observed OTU and the Pielou’s J’ evenness index respectively.

**Figure 2:**
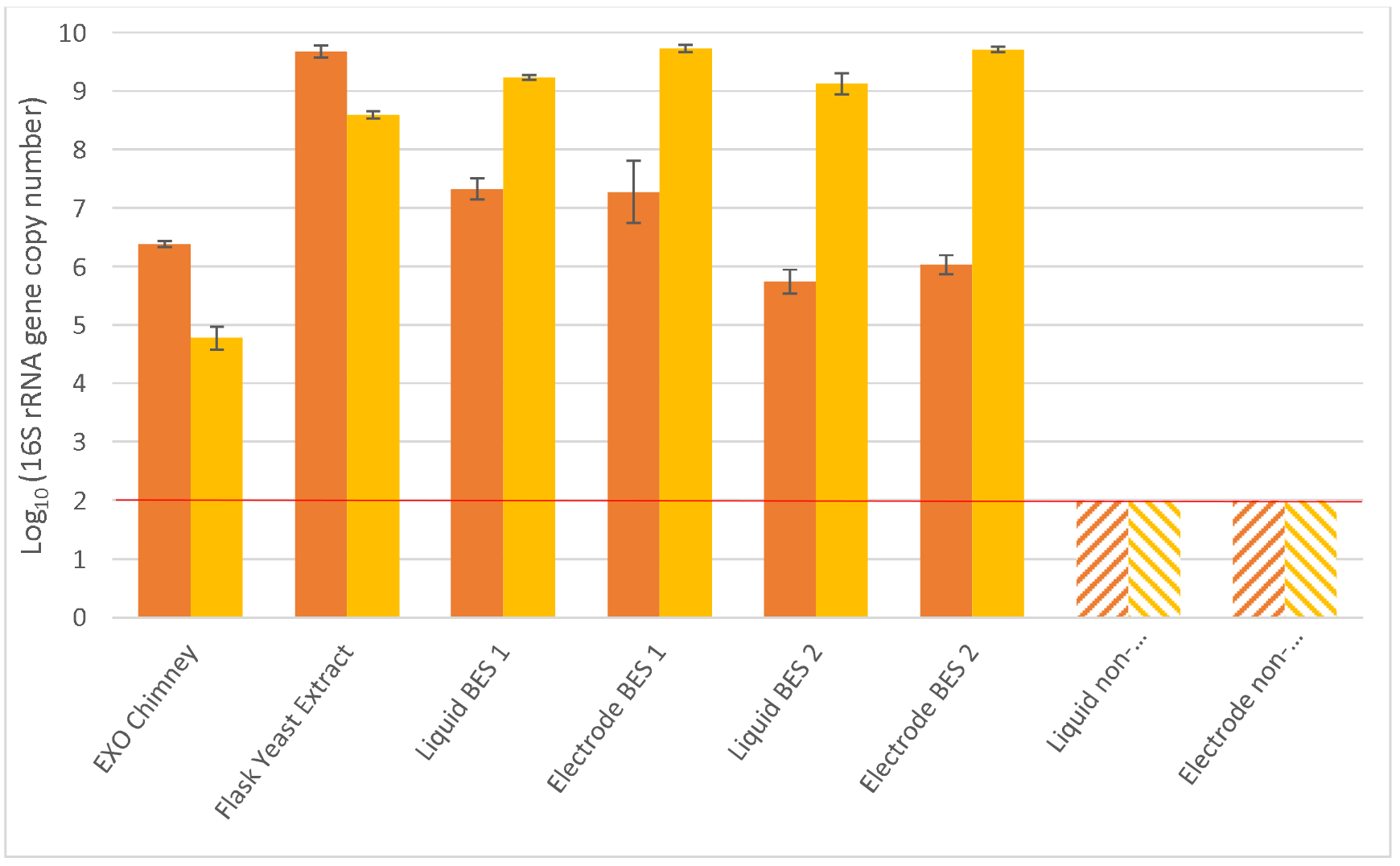
Quantification of 16S rRNA gene copies from Bacteria (orange) or Archaea (yellow) per gram of crushed chimney, per milliliter of liquid or per cm^2^ of working electrode. The red line represents the minimum threshold of sensitivity of the qPCR method. Error bar represent the standard deviation obtained on triplicates.

### 3.2. Enrichment of anode-respiring community in BES from crushed chimney

To assess the diversity of hyperthermophilic exoelectrogenic microbes from deep hydrothermal vent on conductive electrode, a fraction of a crushed chimney from the Rainbow site was used to inoculate the BioElectrochemical System (BES1). To mimic the hydrothermal vent conditions in the BES, a synthetic seawater medium, containing acetate and yeast extract as carbon and energy sources, was used. Indeed, in hydrothermal vent fluid, acetate can be chemically synthesized through Fischer-Tropsch Type (FTT) reaction from H_2_ and CO_2_, both obtained during serpentinization reaction (Schrenk et al. 2013). Acetate can also be biologically produced during fermentative metabolisms. It should be noted that acetate is not fermentable and needs an exogenous electron acceptor to be used as energy source for microorganisms’ growth. On the other hand, YE represent here the organic compounds produced by autotrophic and heterotrophic community of hydrothermal chimney available for the fermentative/respiratory growth of microorganisms.

While in the control experiment (sterile and polarized anode) no notable current (0.01 A/m^2^, figure 3) was observed during the 10 days of experiment, a raise of current density was observed after 6 days of incubation in the BES1 inoculated with crushed chimney. The maximum current density reached 5.9 A/m^2^ at 8.7 days and remained stable for few hours. The 3 mM of acetate consumed during this period as attested by HPLC measurements (data not shown) was in good agreement with the current production assuming a maximum faradic efficiency at 95-100%, as previously obtained in literature (Sengodon and Hays, 2012). After this period the current density decrease progressively probably due to the exhaustion of growth factors present in the medium and necessary to the biofilm metabolic activity. Indeed after 10 days of culture, the renewal of the medium led again to a current increase of up to 6 A/m^2^ at maximum current density. Similarly, the current decreased after a few hours but remained stable in absence of medium renewal.

**Figure 3:**
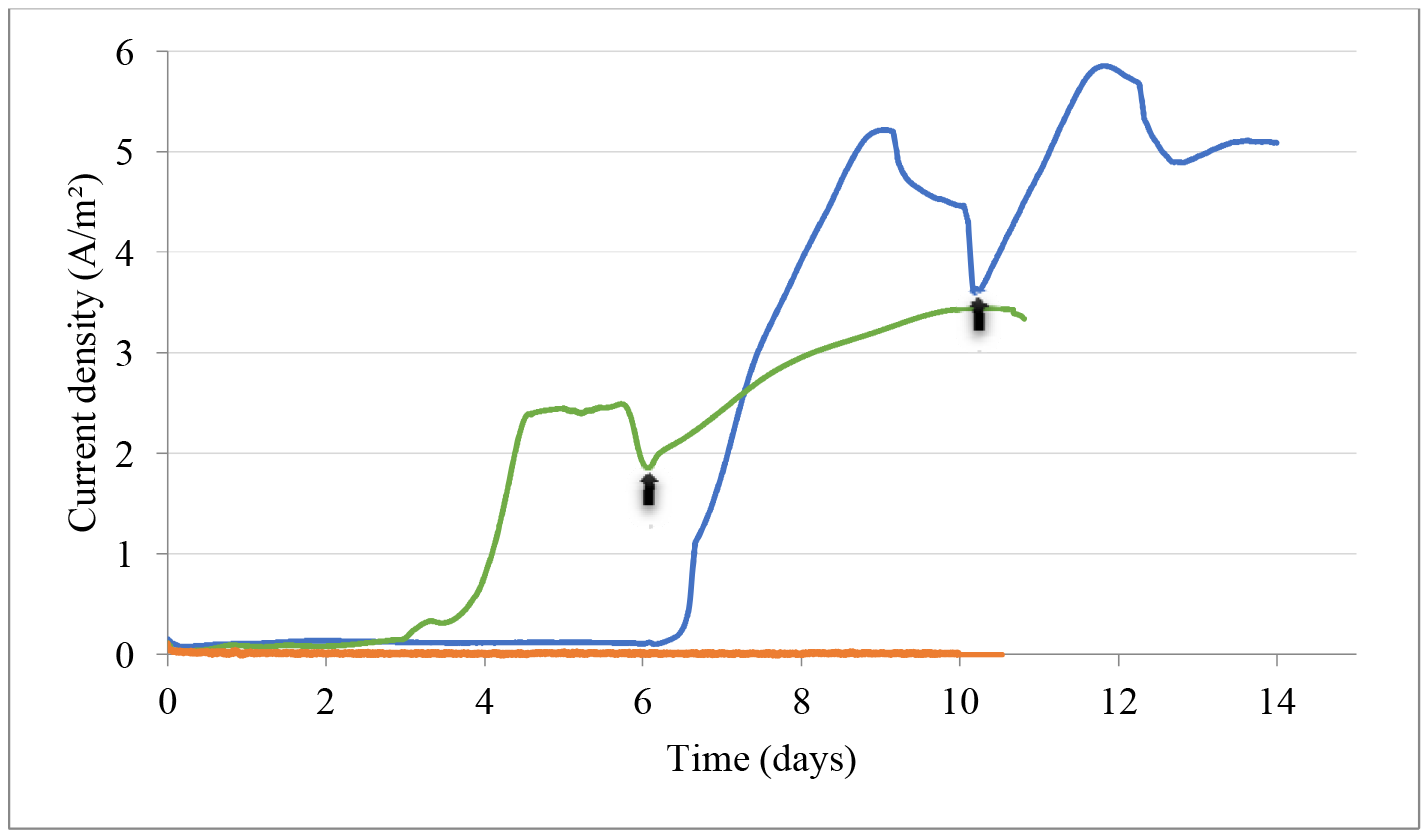
Current monitoring (A/m^2^) of BES1 Enrichment inoculated with crushed hydrothermal chimney (blue line), BES2 Enrichment inoculated with flask subculture (green line) and controls (orange line). Arrows represent the renewal of the liquid media.

Cyclic voltammetries (figure 4) have been performed after inoculation and at the end of experiment in polarized condition to observe the electrochemical profile of ElectroActive Biofilms (EAB) as catalyst of the bioelectrochemical oxidation of acetate. The anode showed the apparition of an oxidation peak with a midpoint potential at +0.06V (vs. SHE) at the end of the experimentation. No oxidation peak was observed for the solution with a new electrode immersed into the medium culture, suggesting EET mechanisms not involving mediators (data not shown). Moreover, the capacitive current (vertical distance between oxidation and reduction waves) increasing between beginning and end of the culture experiments reports an increase of the electrochemical double layer and suggests the presence of a biofilm on the surface of the electrode. Thus, these results suggest the development of EAB with direct EET using acetate as electron donor.

**Figure 4:**
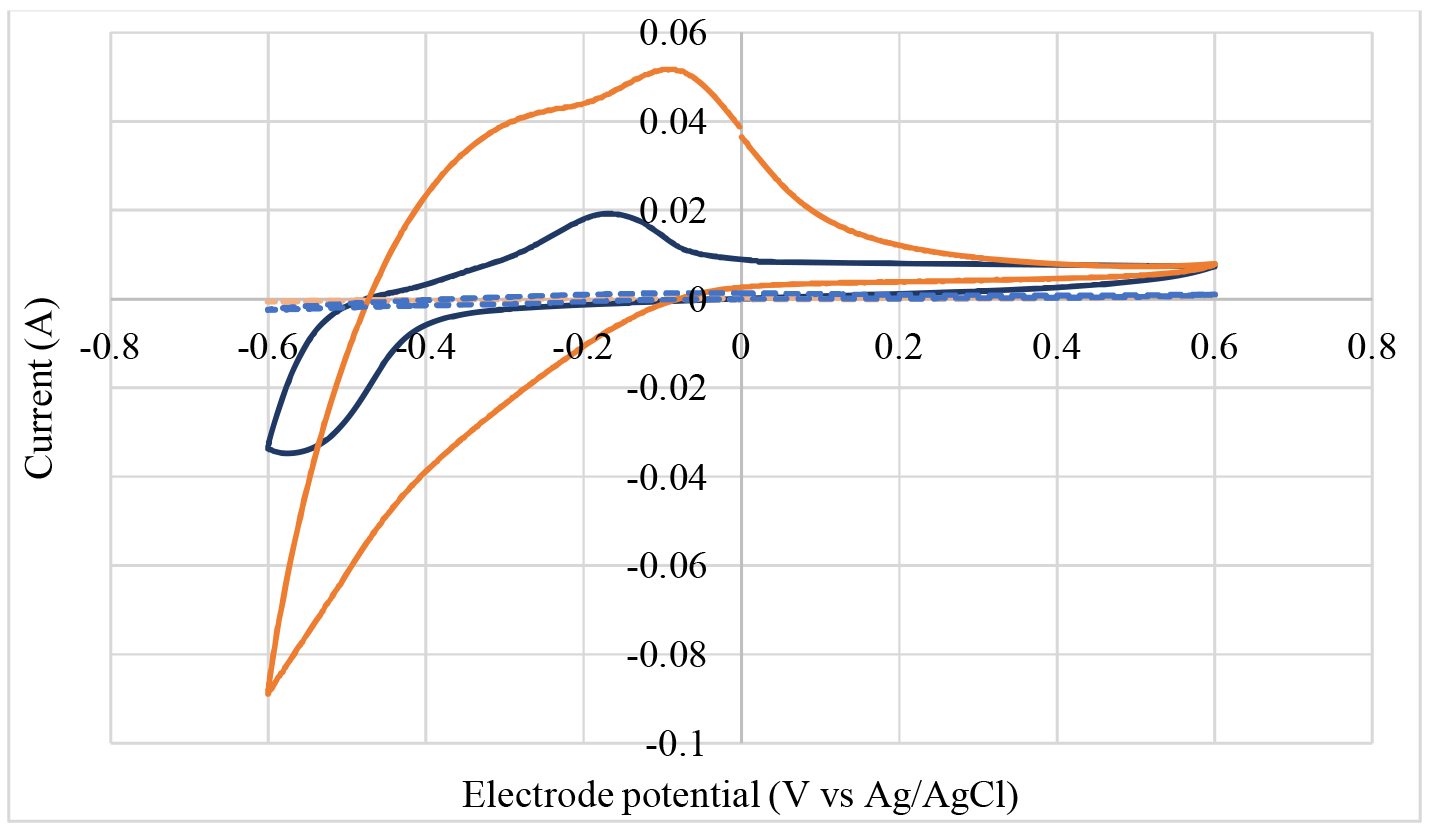
Cyclic Voltammetries between −0,6V and +0,6V at 20mV/s of Working Electrode at initial time (dot lines) and final time (full line) in BES1 (blue line) and BES2 (orange line).

In contrast to the non-polarized culture conditions, microscopic observations over time have revealed the presence of planktonic microorganisms (in the liquid medium) after 9 days of incubation in the polarized electrode culture condition. In addition, the microbial quantification by qPCR of the enrichment in BES1 shows 7.3 ± 0.18 and 9.23 ± 0.04 log of bacterial and archaeal 16S rRNA gene copies per milliliter of liquid media and 7.28 ± 0.53 and 9.73 ± 0.06 log per square centimeter of electrode, respectively. On the contrary, in the non-polarized control, quantification of 16S rRNA copies was under the 2 log of sensitivity threshold of the method, indicating that no significant growth was observed under these conditions in liquid media or non-polarized electrode. Thus, the polarization of the electrode (+ 0.05V vs SHE), which served as an obligatory electrons acceptor to allow the oxidation of acetate or the weak amount of YE, seems essential also for the planktonic microbial growth.

To identify the microbes present on the electrode (Electrode BES1) and in liquid medium (Liquid BES1), a microbial diversity study through sequencing of the hypervariable V4 region of 16S rRNA gene (figure 1) have been performed at the end of the enrichment. 61 OTUs were obtained on Electrode 1 and 58 OTUs in Liquid Media 1, indicating of a loss of biodiversity from environmental chimney (160 OTUs). The sequences obtained are distributed in the different OTUs with however some OTUs more represented than in environmental sample (Equitability at 0.452 on Electrode 1 and 0.331 in Liquid Media 1). This can be explained by our selective conditions of enrichment with high temperature and substrate specificity. Based on average abundance analysis, the microbial diversity of enrichment culture was dominated by *Euryarchaeota* phyla (> 70%), either on electrode or in liquid medium. Interestingly, at the genus level, the dominating archaeal OTUs on electrode were closely related to *Geoglobus* spp. (45.2 %) and *Thermococcus* spp. (25.6%) whereas the planktonic microbial diversity in liquid medium is especially dominated by uncultured *Thermococcus* spp. (64.1%) rather than *Geoglobus* spp. (13.3%). The bacterial diversity on the electrode and in liquid media were mainly assigned to *Thermodesulfatator* spp. (15.6% and 18% respectively), while the remaining were distributed between members of *Bacillaceae* and *Micrococcaceae* families. These specific enrichments of *Archaea* compared to *Bacteria* demonstrate a shift in initial microbial community structure from the crushed chimney, surely driven by the specific BES characteristics (i.e., polarized electrode, hyperthermophilic condition).

Due to the limitation for taxonomic affiliation at species level of the Illumina techniques on 300bp 16S fragments, we performed 16S rDNA clone libraries (figure 5) on ~1300bp 16S sequence. It allowed identifying the *Thermococcus* spp. as closely related to *Thermococcus thioreducens*, *T. coalecens*, *T. barossi*, *T. peptonophilus*, *T. celer* (98% similarities with them) and an unidentified *Thermococcus* species (95% similarity with *Thermococcus thioreducens*). However, because of the high levels of similarity between species of the genus *Thermococcus*, the taxonomic affiliation through the coding gene for 16S rRNA is not sufficient to determine the *Thermococcus* species present in enrichments.

**Figure 5:**
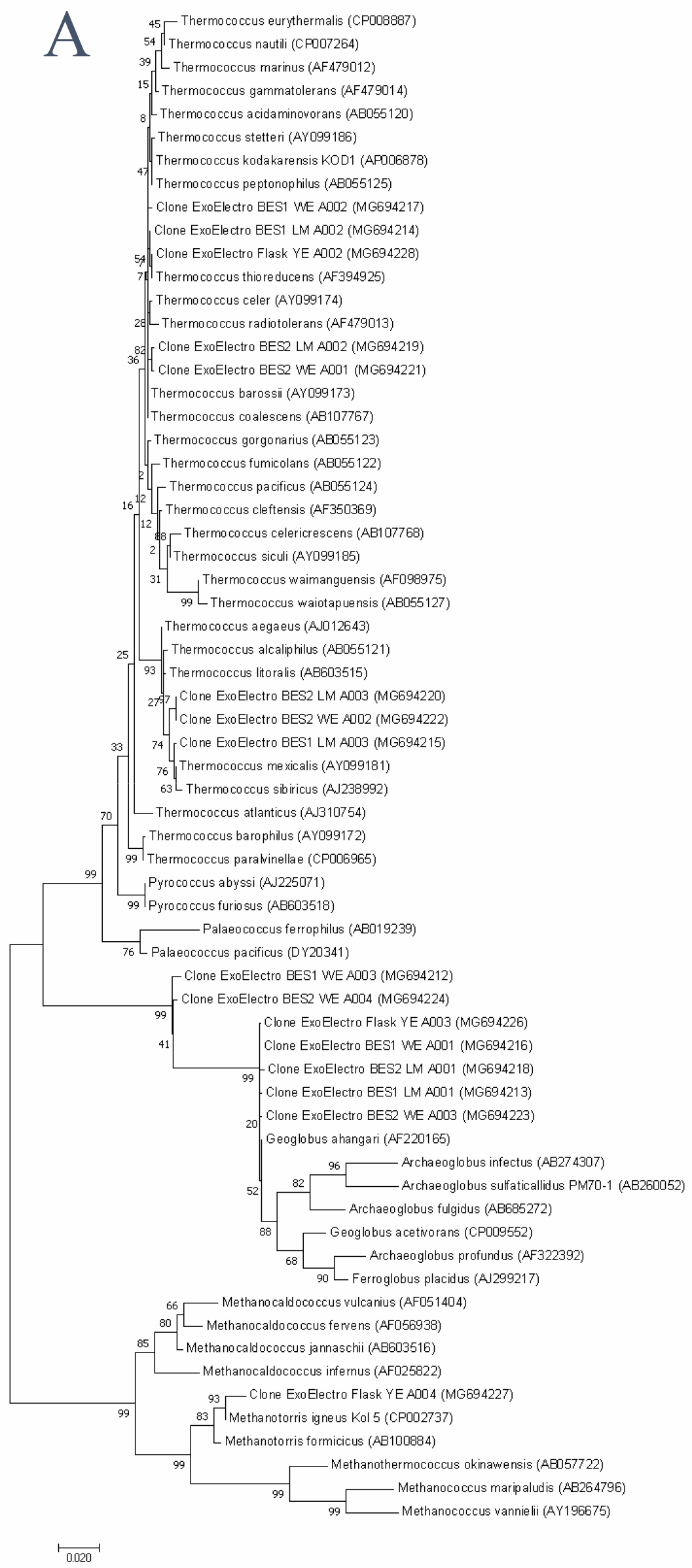
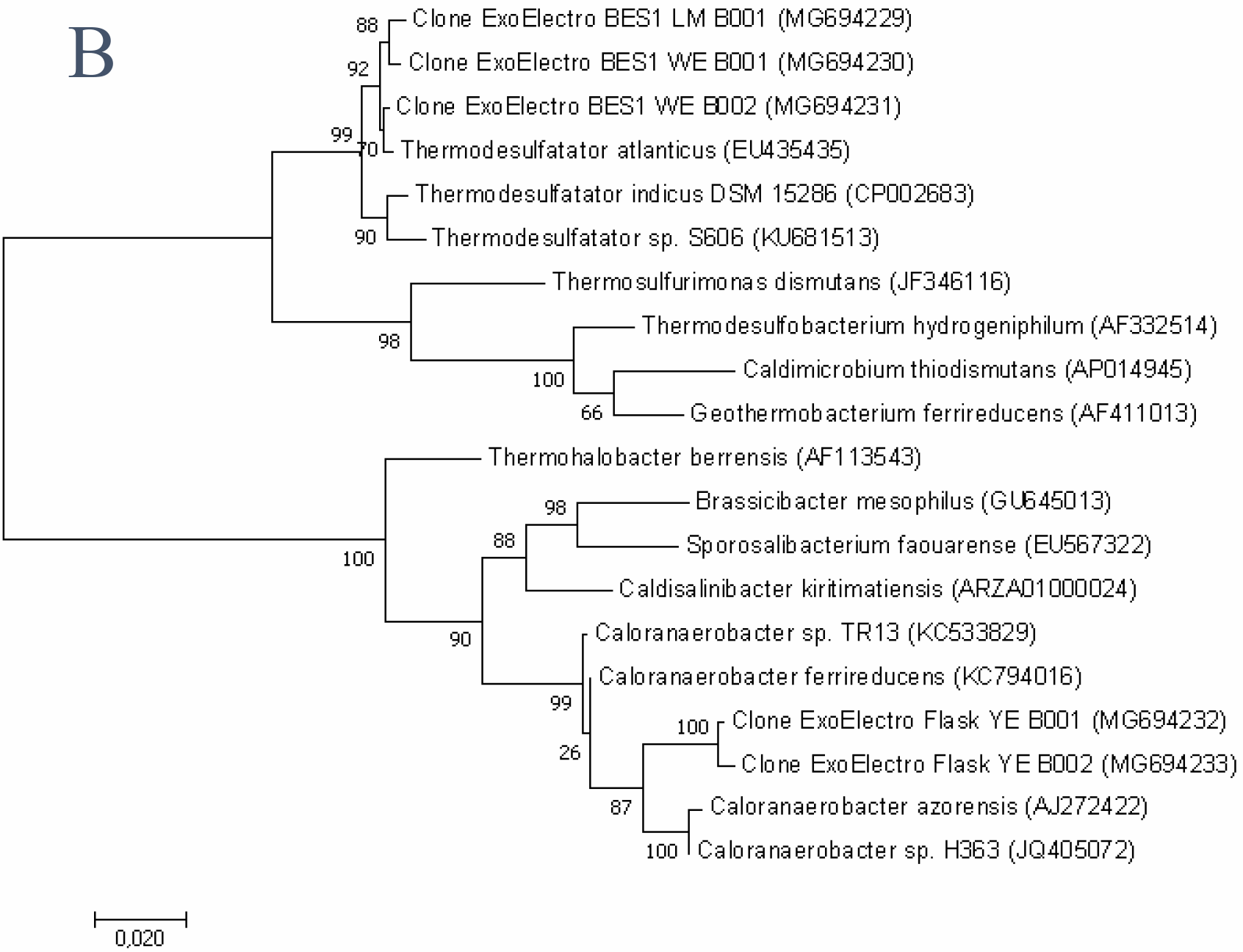
Phylogenetic tree of archaeal (A) and bacterial (B) 16S rRNA clones library from Flask enrichment and Electrode (WE) or liquid media (LM) of BES1 and BES2. Percentages at nodes are bootstrap values based on 500 replications. Scale bar indicated substitutions per site.

However, among the *Geoglobus* spp. the taxonomic affiliation from clone libraries (figure 5) has allowed to identify two species: *Geoglobus ahangari* (99% similarity) and a novel genus or species of the *Archaeoglobaceae* family (MG694212 and MG694224: 95 % of similarity vs *Geoglobus ahangari*). This novel taxon is exclusively found in the electrode area. Interestingly, this affiliation has been possible only through near full 16S rRNA gene sequence (1300 pb) alignment. Indeed, the hypervariable V4 region of the rRNA gene (~290pb), used to the study of microbial diversity by Illumina-based analysis, did not allow to phylogenetically differentiate this novel taxon from *Geoglobus ahangari*. Differences are remarkable at level of hypervariable V6 to V8 regions.

The discovery of *Geoglobus* species enrichment on the electrode is remarkable. The two species of *Geoglobus* described so far (*G. ahangari* and *G. acetivorans*) are known to grow autotrophically using H_2_ or heterotropically using a large number of organic compounds with in both cases soluble or insoluble iron III oxide as a final electrons acceptor (Manzella et al. 2013). More recently, it has been shown that *Geoglobus ahangari* was exoelectrogenic by direct contact when placed in one-chamber microbial electrolysis cells at 80°C. In contrast to our results, the current produced was particularly weak probably due to its use as an axenic culture, and its lack of enzymatic component to resist to oxidative stress on electrode (Yilmazel et al. 2018). Similarly, the type species of *Geoglobus ahangari* tested in our laboratory had a slow and difficult growth on conductive support and was unable to grow after three subcultures under optimal culture conditions. Thus, observing the presence of *Geoglobus* spp. in our system suggests that the latter requires to be cultured in a consortium.

Phylogenetic analysis of 16S rRNA clone libraries has led to the identification of taxon *Thermodesulfatator atlanticus* among the *Thermodesulfatator spp.* previously identified in high throughput sequencing. This bacterium, that have been also isolated from a Rainbow site chimney, is known to be chemolithoautotrophic, sulfate-reducing obligate bacterium that uses H_2_ as electron donor and CO_2_ and peptide as carbon source (Alain et al. 2010). A close related species, *Thermodesulfobacterium commune* has already been identified in EAB on electrode of a MFC (Fu et al. 2015)

### 3.3. Enrichment of hyperthermophilic microorganisms on crystalline iron (III) oxide

In parallel of the first BES previously described, flask enrichments with insoluble metallic electron acceptor were done. Crushed chimney was inoculated in anaerobic mineral medium added with crystalline Fe (III) oxide (α-Fe_2_O_3_: hematite) as electron acceptor (Wahid and Kamalam, 1993), acetate (10 mM) and yeast extract (4 g/l) as energy and carbon source. After 2 days of static incubation at 80°C, the hematite is completely reduced to black compounds (magnetic particles) and the head space of the flask contained 32% of CO_2_, 0.2% of H_2_ and 10% of CH_4_. A production of acetate was also observed in the liquid media, reaching 12 mM at the end of the experiment. The next subcultures reached the black coloration in less than 24 hours with similar concentration of each gas in the headspace of the flasks and same profile of metabolites in the liquid media. The presence of a thin biofilm could be observed on the surface of the agglomerated iron oxide. An abiotic control with a mix of acetate, yeast extract, H_2_, CH_4_ and CO_2_ didn’t show black coloration of iron oxide after one month of incubation at 80°C. These results suggest the development of a community of fermentative organisms producing acetate and electroactive microorganisms able to oxidize organic compounds brought by the YE (peptides, carbohydrates, etc.) and acetate in the culture medium for the reduction of crystalline iron (III) oxide.

The biodiversity enriched in flask was composed of 29 OTUs, and the sequences were relatively concentrated in dominant OTUs (equitability at 0.292). At the family level, the microbial diversity was dominated by bacterial OTUs related to *Clostridiaceae* (73%), and *Thermodesulfobacteriaceae* (3%). The archaeal OTUs represented only 24% of the biodiversity distributed between *Thermococcaceae* (20.5 %), *Methanococcaceae* (3%) identified as *Methanotorris igneus* species and *Archaeoglobaceae* (0.5%). The quantification showed an enrichment of 9 log of bacterial and 7 log of archaeal 16S rRNA gene copies which demonstrated a shift of enrichment toward the preferential bacterial growth in opposition with BES enrichment. Interestingly, the taxonomic affiliation showed that members of the *Clostridiaceae* OTUs were closely related in clone libraries to *Caloranaerobacter ferrireducens* (97% of identity), fermentative microorganism which is known to use crystalline iron (III) reduction as a minor pathway for electron flow while fermenting sugars or amino acids to a mixture of volatile fatty acids (acetate, butyrate) and hydrogen (Zheng et al. 2015). *Archaeoglobaceae* accounted for only 0.5% of the microbial communities in this enrichment and have been closely related from clone libraries analysis to *Geoglobus ahangari* (99% of identity). As mentioned above, *Geoglobus ahangari* is known to reduce amorphous iron (III) oxide and these culture conditions with crystalline form do not seem to favor its growth (Wahid & Kamalan, 1993). Thus, these results could suggest the organization of a kind of trophic chain in our flask enrichment. The new iron reducing species close to *Caloranaerobacter ferrireducens* would have used organic compounds from yeast extract to reduce crystalline iron (III) oxide. The *Thermococcus* would have grown on peptides or carbohydrates from yeast extract or partly produced by *Firmicutes,* and both would have produced H_2_. The methanogens such as *Methanotorris igneus* would then have used the H_2_ to produce methane.

### 3.4. Solid electron acceptor determine the microbial ecology of electroactive enrichment

In this context, a new BioElectrochemical System (BES2) was inoculated with the flask enrichment on YE/acetate to assess the specificity of the microbial community obtained on the electrode. Experimental conditions were exactly the same as for BES1. Contrary to BES1, a lag time of 3 days (figure 3) has been observed before the increase of the current. After 4.5 days the current has reached 2.4 A/m^2^ and stayed stable during about 1 day before decreasing rapidly. Phase contrast microscopic observations during the replacement of the culture medium in BES2 have shown the only presence of irregular coccoid cells. Thereafter, the current density reached a maximum of 3.4 A/m^2^ at 4 days after the renewal of the culture medium. This phase of current production came along with a decrease of acetate as observed in the BES1 (data not shown). Cyclic voltammetry (figure 4) exhibit oxidation peak appearing between initial time and final time with a midpoint potential at +0.053 V (vs. SHE), slightly offset compared to the one obtained with BES1.

The biodiversity indexes between YE enrichment in flask and BES2 showed a loss of richness of species from 29 to 26 OTUs, and equitability from 0.292 to about 0.17, indicating a loss of biodiversity dominated only by a few OTUs. Furthermore, only *Archaea* were detected in BES2, when they were trivial in Flask enrichment. The dominating OTUs (figure 1) were closely related to *Thermococcaceae* uncultured species at 87.2% on electrode and 95.4% in liquid media of total OTUs. The remaining archaeal diversity was composed of *Archaeaoglobaceae* spp. (12.7% and 4.5%), with clones (figure 5a) affiliated to *Geoglobus ahangari* (99% identity) and unknown species belonging to *Archaeoglobaceae* family (95% identity with *Geoglobus ahangari*) similarly found on BES1 electrode. No *Caloranaerobacter* spp. or *Thermodesulfobacteriaceae* spp. growth was observed on the BES2 neither on electrode nor in liquid media. This is supported by the qPCR, which has shown a drastic difference of 4 log between archaeal (9 log) and bacterial (5 log) 16S rRNA copies per milliliter of liquid media and square meter of electrode.

It should be pointed out that the flask YE enrichment did not promote growth of *Archaeoglobales* or *Archaea* in general, whereas they seemed to be re-enriched in the BES2 conditions. These results suggest the specific enrichment on polarized electrode of *Archaea* belonging to *Thermococcales* and *Archeoglobales* in our BES condition. It is noteworthy that the biodiversity indexes showed a loss of biodiversity in BES compared to crushed chimney due to selective condition of our experiment to enrich preferentially the electroactive microorganisms. However, in each BES, the richness and the equitability are more important on polarized electrode than in liquid media, suggesting a specific enrichment of more diverse species and better distribution of microbial population on polarized support. According to the absence of growth in non-polarized electrode and the more important diversity on electrode than in liquid, the microbial diversity found in liquid media would arise from EAB which develop on the electrode. This is supported by the really low H_2_ production in BES (data not shown) - H2 being normally produced by *Thermococcus* sp. during fermentative metabolisms - which could be explained by a lack of carbon and energy source in liquid media. Thus, in absence of possible known metabolism for *Thermococcus* species in liquid, we would suggest *Thermococcus* cells to be in quiescent conditions after it is released from BEA. Its growth would thus have an obligatory dependence on the polarized electrode.

If the presence of *Geoglobus* species is consistent with its known capacity to transfer EET with acetate as electron donor (Yilmazel et al. 2018), *Thermoccoccus* species abundance on polarized electrode is more surprising. *Thermoccocales* are known to share an energetic peptidic or carbohydrate metabolisms, associated necessarily or not with the reduction of elemental sulfur (Bertoldo and Antranikian, 2006). In addition, it was shown that some species have the capacity to growth by using Extracellular Polymeric Substances (EPS) of microbial origin (dextran, pullulan, peptides, etc.) (Legin et al. 1998). So far, no respiration ability on electrode or direct EET mechanism have been reported about this archaeal genera, even if some *Thermococcus* species have been shown to reduce amorphous iron (III) oxide. However, the mechanism is still unclear and could be mediated through electron transfer to humic substances and other extracellular quinones (Lovley et al. 2000, Slobdokin et al. 2001). Furthermore, a recent study has highlighted the possibility of hydroquinone for microbial electron transfer to electrically conductive minerals, especially with pyrite composing hydrothermal chimney, or electrode (Taran, 2017). Nevertheless, no current was obtained with replacement of electrode with spent media, indicating no mediated electron transport in our condition. However, species of *Thermococcus* genus have been reported to produce nanopods or nanotubes by budding of their cell envelope but their function have not been fully understood (Marguet et al. 2013). As suggested by the authors, these archaeal nanopods or nanotubes could be used to expand the metabolic sphere around cells, promote intercellular communication or act in sulfur detoxifying mechanisms (Gorlas et al. 2015). We can therefore suggest a role of these nanotubes in EET as previously described on *Shewanella oneidensis* with the formation of conductive nanofilaments by extensions of the outer membrane. This bacterial EET mechanism involves cytochrome-c on the external membrane, not present in *Thermococcus* genome, allowing the electron transport along the filament to conductive support (Pirbadian et al. 2014). Thus, *Thermococcus* sp. could potentially be exoelectrogenic microorganisms through a still unknown mechanism.

Remarkably, the obligatory presence of both Thermococcus and Geoglobus species found in each BES on the polarized electrode did not seem fortuitous. Assuming that the *Thermococcus* species found in our enrichment are heterotroph, after consumption of YE traces, their only carbon source available would be the EPS or organic compounds produced by *Geoglobus* spp. present on polarized electrode. Then, their fermentative metabolism would lead to the production of acetate, H_2_ and CO_2_ which could then be used by *Geoglobus* species to grow using the electrode as the ultimate electrons acceptor. However, recent data have shown the inability of *Geoglobus* to transfer electrons to an electrode from H_2_ (Yilmazel et al. 2018). Previous studies have shown that some electroactive bacteria are also able to grow syntrophically with other microorganisms via direct interspecies electron transfer (DIET) (Shrestha and Rotaru, 2014). As explained previously, *Geoglobus ahangari* has shown a weak electron transfer capacity when grown in pure culture. The higher current density obtained in our conditions would suggest that the *Geoglobus sp.* and *Thermoccocales* enriched on our polarized electrodes live syntrophically to improve their growth, and subsequently increased the quantity of electrons transferred to the electrode. Thus we would suggest that there is a syntrophic mechanism, with potentially DIET, between *Thermococcus* and *Geoglobus* in deep hydrothermal vents.

## 4. Conclusion

This study is the first to report on the enrichment of electroactive consortium in ex-situ conditions that mimick the conductive chimney of a hydrothermal vent with polarized carbon cloth in anaerobic artificial seawater at 80°C. Moreover, we demonstrate the specificity of enrichment of *Bacteria* on Iron (III) oxide compared to the enrichment of *Archaea* (mainly *Thermococcus sp.* and *Geoglobus sp.*) on BES, where the biodiversity is more conserved. These results comfort the hypothesis of electroactivity as a well-represented metabolism in this type of environment by confirming the presence of exoelectrogenic microorganisms capable of external electron transfer to conductive support.

## Acknowledgements

This work was financially supported by the National Interdisciplinary Program of CNRS “Initiative structurante EC2CO” (MICROBIEN). The authors thank Erwan Roussel (LM2E, IFREMER Brest) for helpful suggestions.

---

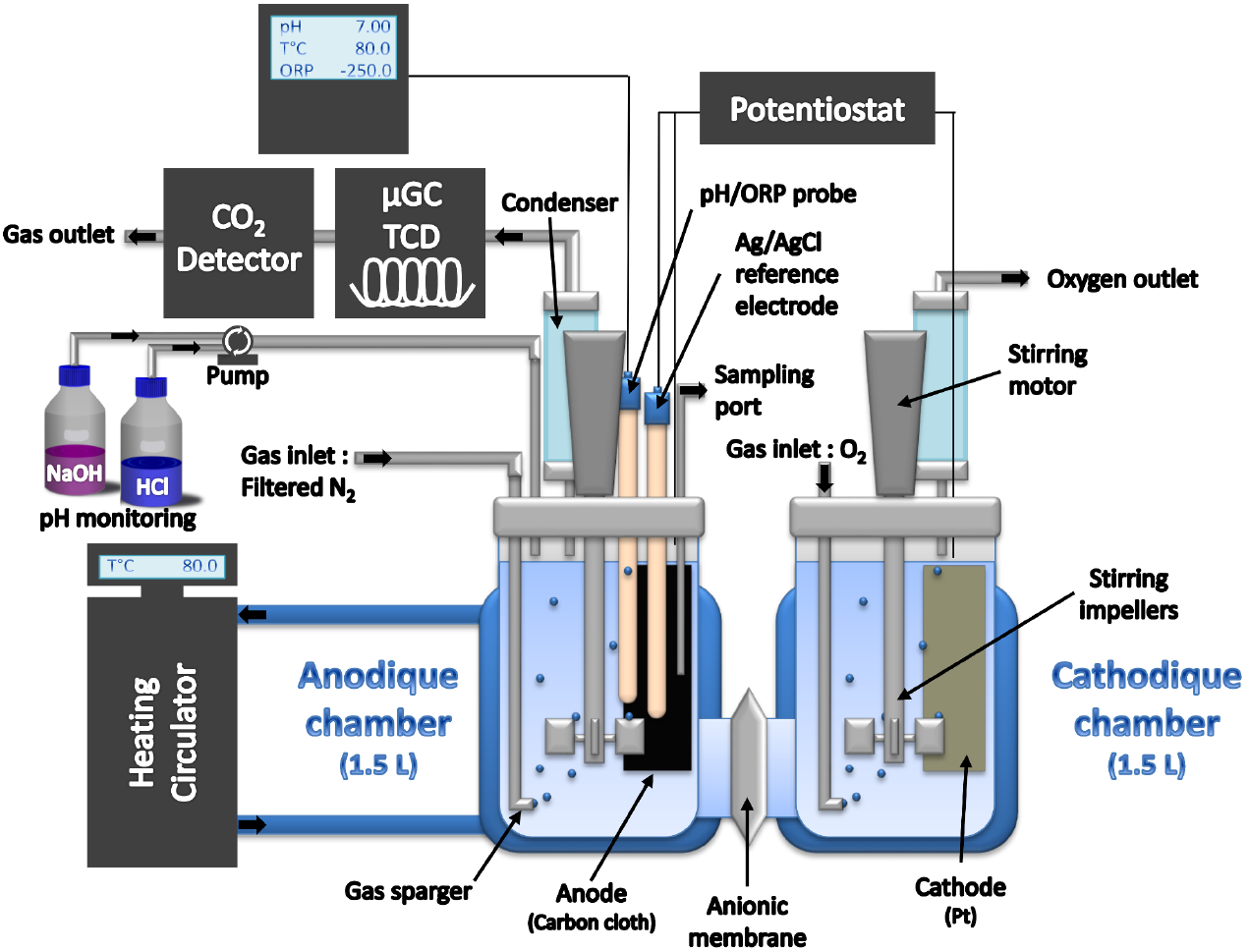

## References

Alain, K., Postec, A., Grinsard, E., Lesongeur, F., Prieur, D., Godfroy, A., 2010. Thermodesulfatator atlanticus sp. nov., a thermophilic, chemolithoautotrophic, sulfate-reducing bacterium isolated from a Mid-Atlantic Ridge hydrothermal vent. Int. J. Syst. Evol. Microbiol. 60, 33–38. https://doi.org/10.1099/ijs.0.009449-0

Allen, R.M., Bennetto, H.P., 1993. Microbial fuel-cells. Appl. Biochem. Biotechnol. 39–40, 27–40. https://doi.org/10.1007/BF02918975

Boileau, C., Auria, R., Davidson, S., Casalot, L., Christen, P., Liebgott, P.-P., Combet-Blanc, Y., 2016. Hydrogen production by the hyperthermophilic bacterium *Thermotoga maritima* part I: effects of sulfured nutriments, with thiosulfate as model, on hydrogen production and growth. Biotechnol. Biofuels 9, 269. https://doi.org/10.1186/s13068-016-0678-8

Busalmen, J.P., Esteve-Núñez, A., Berná, A., Feliu, J.M., 2008. C-Type Cytochromes Wire Electricity-Producing Bacteria to Electrodes. Angew. Chem. 120, 4952–4955. https://doi.org/10.1002/ange.200801310

Cao, H., Wang, Y., Lee, O.O., Zeng, X., Shao, Z., Qian, P.-Y., 2014. Microbial Sulfur Cycle in Two Hydrothermal Chimneys on the Southwest Indian Ridge. mBio 5, e00980-13–e00980-13. https://doi.org/10.1128/mBio.00980-13

Cerqueira, T., Pinho, D., Froufe, H., Santos, R.S., Bettencourt, R., Egas, C., 2017. Sediment Microbial Diversity of Three Deep-Sea Hydrothermal Vents Southwest of the Azores. Microb. Ecol. 74, 332–349. https://doi.org/10.1007/s00248-017-0943-9

Dick, G.J., Anantharaman, K., Baker, B.J., Li, M., Reed, D.C., Sheik, C.S., 2013. The microbiology of deep-sea hydrothermal vent plumes: ecological and biogeographic linkages to seafloor and water column habitats. Front. Microbiol. 4. https://doi.org/10.3389/fmicb.2013.00124

Doyle, L.E., Marsili, E., 2015. Methods for enrichment of novel electrochemically-active microorganisms. Bioresour. Technol. https://doi.org/10.1016/j.biortech.2015.07.025

Flores, G.E., Campbell, J.H., Kirshtein, J.D., Meneghin, J., Podar, M., Steinberg, J.I., Seewald, J.S., Tivey, M.K., Voytek, M.A., Yang, Z.K., Reysenbach, A.-L., 2011. Microbial community structure of hydrothermal deposits from geochemically different vent fields along the Mid-Atlantic Ridge. Environ. Microbiol. 13, 2158–2171. https://doi.org/10.1111/j.1462-2920.2011.02463.x

Fu, Q., Fukushima, N., Maeda, H., Sato, K., Kobayashi, H., 2015. Bioelectrochemical analysis of a hyperthermophilic microbial fuel cell generating electricity at temperatures above 80 °C. Biosci. Biotechnol. Biochem. 0, 1–7. https://doi.org/10.1080/09168451.2015.1015952

Fu, Q., Kobayashi, H., Kawaguchi, H., Wakayama, T., Maeda, H., Sato, K., 2013. A Thermophilic Gram-Negative Nitrate-Reducing Bacterium, Calditerrivibrio nitroreducens, Exhibiting Electricity Generation Capability. Environ. Sci. Technol. 47, 12583–12590. https://doi.org/10.1021/es402749f

Girguis, P.R., Holden, J.F., 2012. On the Potential for Bioenergy and Biofuels from Hydrothermal Vent Microbes. Oceanography.

Godfroy, A., 2005. EXOMAR cruise, RV L’Atalante [WWW Document]. URL http://dx.doi.org/10.17600/5010100

Gorby, Y.A., Yanina, S., McLean, J.S., Rosso, K.M., Moyles, D., Dohnalkova, A., Beveridge, T.J., Chang, I.S., Kim, B.H., Kim, K.S., Culley, D.E., Reed, S.B., Romine, M.F., Saffarini, D.A., Hill, E.A., Shi, L., Elias, D.A., Kennedy, D.W., Pinchuk, G., Watanabe, K., Ishii, S., Logan, B., Nealson, K.H., Fredrickson, J.K., 2006. Electrically conductive bacterial nanowires produced by Shewanella oneidensis strain MR-1 and other microorganisms. Proc. Natl. Acad. Sci. 103, 11358–11363. https://doi.org/10.1073/pnas.0604517103

Ha, P.T., Lee, T.K., Rittmann, B.E., Park, J., Chang, I.S., 2012. Treatment of Alcohol Distillery Wastewater Using a Bacteroidetes-Dominant Thermophilic Microbial Fuel Cell. Environ. Sci. Technol. 46, 3022–3030. https://doi.org/10.1021/es203861v

Hinks, J., Zhou, M., Dolfing, J., 2017. Microbial Electron Transport in the Deep Subsurface, in: Chénard, C., Lauro, F.M. (Eds.), Microbial Ecology of Extreme Environments. Springer International Publishing, Cham, pp. 81–102.

Hogarth, M.P., 1995. The development of the direct methanol fuel cell. (Ph.D.). University of Newcastle upon Tyne.

Huber, J.A., Cantin, H.V., Huse, S.M., Mark Welch, D.B., Sogin, M.L., Butterfield, D.A., 2010. Isolated communities of *Epsilonproteobacteria* in hydrothermal vent fluids of the Mariana Arc seamounts: *Epsilonproteobacteria* in vents of the Mariana Arc. FEMS Microbiol. Ecol. no-no. https://doi.org/10.1111/j.1574-6941.2010.00910.x

J. McCormick, A., Bombelli, P., M. Scott, A., J. Philips, A., G. Smith, A., C. Fisher, A., J. Howe, C., 2011. Photosynthetic biofilms in pure culture harness solar energy in a mediatorless bio-photovoltaic cell (BPV) system. Energy Environ. Sci. 4, 4699–4709. https://doi.org/10.1039/C1EE01965A

Jong, B.C., Kim, B.H., Chang, I.S., Liew, P.W.Y., Choo, Y.F., Kang, G.S., 2006. Enrichment, Performance, and Microbial Diversity of a Thermophilic Mediatorless Microbial Fuel Cell. Environ. Sci. Technol. 40, 6449–6454. https://doi.org/10.1021/es0613512

Koch, C., Harnisch, F., 2016. Is there a Specific Ecological Niche for Electroactive Microorganisms? ChemElectroChem 3, 1282–1295. https://doi.org/10.1002/celc.201600079

Konn, C., Charlou, J.L., Holm, N.G., Mousis, O., 2015. The Production of Methane, Hydrogen, and Organic Compounds in Ultramafic-Hosted Hydrothermal Vents of the Mid-Atlantic Ridge. Astrobiology 15, 381–399. https://doi.org/10.1089/ast.2014.1198

Kristall, B., Kelley, D.S., Hannington, M.D., Delaney, J.R., 2006. Growth history of a diffusely venting sulfide structure from the Juan de Fuca Ridge: A petrological and geochemical study: DIFFUSELY VENTING SULFIDE STRUCTURE. Geochem. Geophys. Geosystems 7, n/a–n/a. https://doi.org/10.1029/2005GC001166

Kumar, A., Hsu, L.H.-H., Kavanagh, P., Barrière, F., Lens, P.N.L., Lapinsonnière, L., V, J.H.L., Schröder, U., Jiang, X., Leech, D., 2017. The ins and outs of microorganism–electrode electron transfer reactions. Nat. Rev. Chem. 1, 24. https://doi.org/10.1038/s41570-017-0024

Legin, E., Copinet, A., Duchiron, F., 1998. Production of thermostable amylolytic enzymes by *Thermococcus hydrothermalis*. Biotechnol. Lett. 20, 363–367. https://doi.org/10.1023/A:1005375213196

Logan, B.E., Hamelers, B., Rozendal, R., Schröder, U., Keller, J., Freguia, S., Aelterman, P., Verstraete, W., Rabaey, K., 2006. Microbial fuel cells□: Methodology and technology. Environ. Sci. Technol. 40, 5181–5192.

Lusk, B.G., Khan, Q.F., Parameswaran, P., Hameed, A., Ali, N., Rittmann, B.E., Torres, C.I., 2015. Characterization of Electrical Current-Generation Capabilities from Thermophilic Bacterium *Thermoanaerobacter pseudethanolicus* Using Xylose, Glucose, Cellobiose, or Acetate with Fixed Anode Potentials. Environ. Sci. Technol. 49, 14725–14731. https://doi.org/10.1021/acs.est.5b04036

Marshall, C.W., May, H.D., 2009. Electrochemical evidence of direct electrode reduction by a thermophilic Gram-positive bacterium, *Thermincola ferriacetica*. Energy Environ. Sci. 2, 699. https://doi.org/10.1039/b823237g

Marsili, E., Baron, D.B., Shikhare, I.D., Coursolle, D., Gralnick, J.A., Bond, D.R., 2008. *Shewanella* secretes flavins that mediate extracellular electron transfer. Proc. Natl. Acad. Sci. 105, 3968–3973. https://doi.org/10.1073/pnas.0710525105

Mathis, B.J., Marshall, C.W., Milliken, C.E., Makkar, R.S., Creager, S.E., May, H.D., 2007. Electricity generation by thermophilic microorganisms from marine sediment. Appl. Microbiol. Biotechnol. 78, 147–155. https://doi.org/10.1007/s00253-007-1266-4

Miroshnichenko, M.L., Bonch-Osmolovskaya, E.A., 2006. Recent developments in the thermophilic microbiology of deep-sea hydrothermal vents. Extremophiles 10, 85–96. https://doi.org/10.1007/s00792-005-0489-5

Reguera, G., Nevin, K.P., Nicoll, J.S., Covalla, S.F., Woodard, T.L., Lovley, D.R., 2006. Biofilm and Nanowire Production Leads to Increased Current in *Geobacter sulfurreducens* Fuel Cells. Appl. Environ. Microbiol. 72, 7345–7348. https://doi.org/10.1128/AEM.01444-06

Schrenk, M.O., Brazelton, W.J., Lang, S.Q., 2013. Serpentinization, Carbon, and Deep Life. Rev. Mineral. Geochem. 75, 575–606. https://doi.org/10.2138/rmg.2013.75.18

Schröder, U., Harnisch, F., Angenent, L.T., 2015. Microbial electrochemistry and technology: terminology and classification. Energy Env. Sci 8, 513–519. https://doi.org/10.1039/C4EE03359K

Sekar, N., Umasankar, Y., P. Ramasamy, R., 2014. Photocurrent generation by immobilized cyanobacteria via direct electron transport in photo-bioelectrochemical cells. Phys. Chem. Chem. Phys. 16, 7862–7871. https://doi.org/10.1039/C4CP00494A

Sekar, N., Wu, C.-H., Adams, M.W.W., Ramasamy, R.P., 2017. Electricity generation by Pyrococcus furiosus in microbial fuel cells operated at 90°C. Biotechnol. Bioeng. 114, 1419–1427. https://doi.org/10.1002/bit.26271

Sengodon, P., Hays, D.B., 2012. Microbial Fuel Cells. Future Fuel Technologies, National Petroleum Council (NPC) Study

Shehab, N.A., Ortiz-Medina, J.F., Katuri, K.P., Hari, A.R., Amy, G., Logan, B.E., Saikaly, P.E., 2017. Enrichment of extremophilic exoelectrogens in microbial electrolysis cells using Red Sea brine pools as inocula. Bioresour. Technol. 239, 82–86. https://doi.org/10.1016/j.biortech.2017.04.122

Shi, L., Dong, H., Reguera, G., Beyenal, H., Lu, A., Liu, J., Yu, H.-Q., Fredrickson, J.K., 2016. Extracellular electron transfer mechanisms between microorganisms and minerals. Nat. Rev. Microbiol. 14, 651–662. https://doi.org/10.1038/nrmicro.2016.93

Shi, L., Squier, T.C., Zachara, J.M., Fredrickson, J.K., 2007. Respiration of metal (hydr)oxides by *Shewanella* and *Geobacter*: a key role for multihaem c-type cytochromes. Mol. Microbiol. 65, 12–20. https://doi.org/10.1111/j.1365-2958.2007.05783.x

Shrestha, P.M., Rotaru, A.-E., 2014. Plugging in or going wireless: strategies for interspecies electron transfer. Front. Microbiol. 5, 237. https://doi.org/10.3389/fmicb.2014.00237

Stoddard, S.F., Smith, B.J., Hein, R., Roller, B.R.K., Schmidt, T.M., 2015. rrnDB: improved tools for interpreting rRNA gene abundance in bacteria and archaea and a new foundation for future development. Nucleic Acids Res. 43, D593–598. https://doi.org/10.1093/nar/gku1201

Uzarraga, R., Auria, R., Davidson, S., Navarro, D., Combet-Blanc, Y., 2011. New Cultural Approaches for Microaerophilic Hyperthermophiles. Curr. Microbiol. 62, 346–350. https://doi.org/10.1007/s00284-010-9712-4

Vetriani, C., Voordeckers, J.W., Crespo-Medina, M., O’Brien, C.E., Giovannelli, D., Lutz, R.A., 2014. Deep-sea hydrothermal vent *Epsilonproteobacteria* encode a conserved and widespread nitrate reduction pathway (Nap). ISME J. 8, 1510–1521. https://doi.org/10.1038/ismej.2013.246

Wirth, R., 2017. Colonization of Black Smokers by Hyperthermophilic Microorganisms. Trends Microbiol. 25, 92–99. https://doi.org/10.1016/j.tim.2016.11.002

Wrighton, K.C., Agbo, P., Warnecke, F., Weber, K.A., Brodie, E.L., DeSantis, T.Z., Hugenholtz, P., Andersen, G.L., Coates, J.D., 2008. A novel ecological role of the Firmicutes identified in thermophilic microbial fuel cells. ISME J. 2, 1146–1156. https://doi.org/10.1038/ismej.2008.48

Wrighton, K.C., Thrash, J.C., Melnyk, R.A., Bigi, J.P., Byrne-Bailey, K.G., Remis, J.P., Schichnes, D., Auer, M., Chang, C.J., Coates, J.D., 2011. Evidence for Direct Electron Transfer by a Gram-Positive Bacterium Isolated from a Microbial Fuel Cell. Appl. Environ. Microbiol. 77, 7633–7639. https://doi.org/10.1128/AEM.05365-11

Yamamoto, M., Nakamura, R., Kasaya, T., Kumagai, H., Suzuki, K., Takai, K., 2017. Spontaneous and Widespread Electricity Generation in Natural Deep-Sea Hydrothermal Fields. Angew. Chem. Int. Ed. 56, 5725–5728. https://doi.org/10.1002/anie.201701768

Yilmazel, Y.D., Zhu, X., Kim, K.-Y., Holmes, D.E., Logan, B.E., 2018. Electrical current generation in microbial electrolysis cells by hyperthermophilic archaea *Ferroglobus placidus* and *Geoglobus ahangari*. Bioelectrochemistry 119, 142–149. https://doi.org/10.1016/j.bioelechem.2017.09.012

Zhang, L., Kang, M., Xu, J., Xu, J., Shuai, Y., Zhou, X., Yang, Z., Ma, K., 2016. Bacterial and archaeal communities in the deep-sea sediments of inactive hydrothermal vents in the Southwest India Ridge. Sci. Rep. 6. https://doi.org/10.1038/srep25982

